# Myo-inositol and total NAA in the hippocampus are linked to CSF tau pathology in cognitively normal older adults

**DOI:** 10.1101/2024.08.09.607353

**Authors:** JA Noche, C Vanderlip, S Wright, L. Sordo, E Head, CEL Stark

## Abstract

**INTRODUCTION:** Understanding relationships between *in vivo* neurometabolic changes and Alzheimer’s disease (AD) pathology in the hippocampus, a region vulnerable to early changes in AD, will support early diagnosis.

**METHODS:** Two studies using ^1^H-MRS examined concentrations of myo-inositol (MI), total creatine (tCr) and total NAA (tNAA) in the hippocampus. The first study compared hippocampal metabolite concentrations in healthy young and older adults and the second study assessed relationships between hippocampal metabolites and cerebrospinal fluid (CSF) measurements of Aβ42, phosphotau 181 (pTau181), and total tau (t-Tau) while adjusting for demographic covariates and spectral characteristics (linewidth, signal-to-noise ratio) in a separate group of older adults ranging from cognitively normal (CN) to AD-dementia.

**RESULTS:** Hippocampal MI, but not tCr or tNAA, was increased in cognitively normal older versus young adults. Within the second older adult group, MI and tNAA, but not tCr, were linked to increases in CSF pTau181 and t-Tau, but not Aβ42.

**DISCUSSION:** Tau deposition in cognitively normal individuals is associated with biochemical changes related to glial reactivity and neural integrity in the hippocampus.

## Introduction

Pathological hallmarks of Alzheimer’s disease (AD) such as the extracellular accumulation of beta-amyloid (Aβ) plaques and the intraneuronal accumulation of neurofibrillary tangles comprised of aggregated hyperphosphorylated tau begin decades before individuals exhibit cognitive decline^1^. Improving our understanding of how these pathologies alter brain function can help to identify novel therapeutic targets and enable the development of non-invasive biomarkers of AD for early and accurate diagnosis. Proton magnetic resonance spectroscopy (^1^H-MRS) is a non-invasive, *in vivo* technique within magnetic resonance imaging (MRI) that has proven useful in measuring neurometabolites known to become altered in aging and AD. Common, readily-assessed metabolites include the neuronal marker n-acetyl aspartate (NAA), an amino acid mainly synthesized and stored in neurons^2^ that declines with age, cognitive performance, and in neurodegenerative disorders including AD^3^. Myo-inositol (MI) is an osmolyte primarily found in glia that is a putative marker of glial reactivity and neuroinflammation^4,5^. Importantly, neuroinflammation is viewed as a hallmark of AD, and increases to MI may be a result of reactive gliosis in this disease^6–8^. Creatine (Cr) and phosphocreatine (PCr) are also readily quantified and are involved in energy metabolism. Either Cr alone or its total composite (tCr) is commonly used as an internal reference with the longstanding assumption that Cr is stable over the lifespan and within various neuropathologies. However, recent studies suggest this may not be the case^9,10^. Nonetheless, numerous past studies of AD that evaluated metabolite ratios from within the posterior cingulate cortex (PCC) demonstrate that MI and NAA normalized to Cr are altered in AD. For example, PCC MI/Cr ratios are increased in individuals with Mild Cognitive Impairment (MCI) compared to age-matched controls^11^ and MI/Cr is elevated in APOE ε4 carriers before detectable amyloid pathology by CSF Aβ42^12^. Furthermore, several studies report that NAA/Cr is predictive of progression from MCI to dementia^13–15^. However, as these detailed characterizations of biochemical changes in response to AD pathologic burden have focused on the PCC, there are limited reports on similar changes taking place in the hippocampus likely due to the technical challenges in acquiring reliable ^1^H-MRS data from this subcortical region compared to neocortical areas^16^.

A critical region for memory, the hippocampus, is one of the earliest areas affected in AD with hippocampal atrophy, hyperactivity, and associated memory decline, which are considered as some of the most prominent early features of AD prior to clinical diagnosis^17–19^. Furthermore, the hippocampus is an early site for neurofibrillary tangles (NFT) accumulation (Braak stage II)^20^. Animal studies also demonstrate that the hippocampus is vulnerable to early, heightened sensitivity to glial reactivity and neuroinflammation in aging^21^. However, such characterizations of regional neuroinflammation in humans are limited, despite sustained systemic neuroinflammation considered a prominent early contributor to AD pathogenesis^22^.

The recent availability of advanced, open-access MRS software as well as published consensus recommendations for fitting, quantification, and reporting enable robustness in spectral quality assessments, enhancements in signal to noise, and tissue-corrected absolute metabolite quantitation^23^. Therefore, in this study, we sought to determine the effects of aging and AD pathologies on neurometabolites using these modern techniques in ^1^H-MRS processing and analysis within the hippocampus in cognitively normal older adults early along the AD continuum.

We quantified fully tissue-corrected, water referenced concentrations of MI, tCr, and total NAA (tNAA) using point-resolved spectroscopy (PRESS) from the hippocampus of young and older adults to understand typical age-related changes to these AD-relevant metabolites. Then in a separate group of older adults that also had available measures of cerebrospinal fluid (CSF) measures of amyloid (Aβ42), phosphorylated tau 181 (ptau181), and total tau (t-Tau)^24^, we evaluated how hippocampal neurometabolites are altered as a function of global levels of amyloid and tau pathology. We hypothesized that early AD biomarker increases are associated with biochemical alterations related to neuroinflammation, bioenergetics, and neuronal integrity of the hippocampus, even within cognitively normal adults.

## Materials and Methods

### Subjects

Study 1: young adults ages 18-25 and healthy older adults ages 60-85 were recruited from the greater Orange County community. A subset of older adults was additionally recruited in conjunction with the UCI Alzheimer’s Disease Research Center (UCI ADRC), a National Institute on Aging (NIA)-funded research center that conducts prospective longitudinal studies to investigate brain changes in normal aging, MCI, and AD. Individuals recruited through the UCI ADRC were previously assessed by an ADRC study clinician and identified as either cognitively normal with or mild cognitive impairment (MCI) within one year of the MRI scan. Those with MCI were eligible to participate in the study to preliminarily evaluate the progression of neurochemical changes that may exist in prodromal AD. All participants underwent a single MRI session, where the community-recruited participants additionally underwent brief cognitive testing prior to their MRI scan.

Study 2: a separate group of ADRC participants were recruited for participation in a single MRI session comprised of primarily cognitively normal older adults. Like in Study 1, a small subset of participants was included that had non-normal cognition, including questionable cognitive impairment (QCI), MCI, or AD dementia, at their most recent ADRC annual testing session within one year of the MRI session to for early assessments of the progression of neurochemical changes that may exist along different syndrome classifications (Table 1).

**Table 1.** Demographics for each group of MRS subjects from Study 1 and Study 2. Age and MOCA Total Scores are shown as the mean (SD). CN=cognitively normal older adults; QCI=questionable cognitive impairment; MCI=mild cognitive impairment; Dem=AD dementia.

|  | Study 1:<br>Young | Study 1:<br>CN | Study 1:<br>MCI | Study 2:<br>CN | Study 2:<br>QCI | Study 2:<br>MCI | Study 2:<br>Dem |
| --- | --- | --- | --- | --- | --- | --- | --- |
| N | 22 | 15 | 3 | 29 | 3 | 2 | 1 |
| Sex (Female) | 11 | 7 | 2 | 17 | 2 | 1 | 0 |
| Age | 21.14<br>(3.5) | 73.07<br>(4.51) | 67.7<br>(7.02) | 74.21<br>(5.78) | 76.33<br>(7.09) | 78.5<br>(3.53) | 60 |
| MOCA Total<br>Score | 27.2 (1.9) | 27.17<br>(2.7) | 26.0<br>(2.0) | 27.76<br>(1.68) | 24 (1.73) | 22<br>(2.82) | 8 |

For the participants from Study 2, a subset of participants (n=21) had available CSF samples (Table 2). Immunoassays of CSF samples closest to the MRI scan visit were performed at the UCI ADRC Neuropathology Core using the Roche Elecsys® system for measuring quantities of Aβ42 (Elecsys® beta-Amyloid (1-42) CSF II), ptau181 (Elecsys® Phospho-Tau (181P) CSF), and t-Tau (Elecsys® Total-Tau CSF), biomarkers of amyloid pathology, tau pathology, and neurodegeneration, respectively. The delay between CSF sample collection and subsequent MRI scanning session varied across participants (3 ± 2 years). The estimates for Aβ42 were then adjusted by a factor of 0.78 recommended by Roche specifically for this study sample. CSF measures were analyzed as continuous measurements rather than defining binary categorizations of amyloid/tau/neurodegeneration (A/T/N) status. However, based on existing ranges of abnormal CSF biomarker thresholds^25^, our cognitively normal CSF participants are likely considered in the early stages of the AD continuum^26^.

**Table 2.** Study 2 participant characteristics with available CSF immunoassay measurements of Aβ42, ptau181, and t-Tau pg/mL mean (SD).

| Study 2 CSF Subjects | CN | QCI | MCI | Dem |
| --- | --- | --- | --- | --- |
| N | 18 | 1 | 1 | 1 |
| Sex (Female) | 11 | 1 | 1 | 0 |
| Age | 75.17 (6.17) | 75 | 76 | 60 |
| MOCA Total Score | 27.78(1.73) | 25 | 24 | 8 |
| CSF Aβ42 pg/mL | 471.09 (181.15) | 257.166 | 212.238 | 193.44 |
| CSF ptau181 pg/mL | 16.97 (5.58) | 11.78 | 24.37 | 24.15 |
| CSF t-Tau | 182.74 (54.78) | 140.9 | 238.4 | 236.4 |

All participants provided written informed consent and were compensated for their participation in MRI, neuropsychological testing and for providing CSF in compliance with the University of California, Irvine (UCI) Institutional Review Board (IRB). Participants were screened for neurological disease or psychiatric illness, spoke fluent English, were right-handed, had normal or corrected-to-normal vision, and had no contraindications for MRI.

### Single-Voxel 1H MR Spectroscopy

All MR imaging and spectroscopy data were collected using a ^1^H 32-channel transmit/receive head coil on a 3T Siemens MAGNETOM Prisma scanner (Siemens Healthineers, Erlangen, Germany) at the UCI Facility for Imaging and Brain Research (FIBRE). Scanning sessions for both studies lasted approximately one hour. T1-weighted (T1w) whole-brain anatomical images were acquired using a 3D magnetization-prepared rapid gradient echo (MP-RAGE) for Study 1 with the following parameters: resolution = 0.8 mm isotropic; repetition time (TR) and echo time (TE) = 2300/2.38ms; field of view = 240 x 320 x 320; flip angle = 8°. Study 2 used the Alzheimer’s Disease Neuroimaging Initiative 3 (ADNI-3) sequence parameters: resolution = 1.0 mm isotropic; TR/TE = 2300/2.98ms; flip angle = 9°.

The single ^1^H-MRS voxel of interest (VOI) was aligned to the long axis of the right hippocampus containing mainly the anterior hippocampus using either the T1w image or a multi-slice scout image as an anatomical reference (Figure 1). Both studies used the vendor-supplied ^1^H-MRS PRESS sequence. Study 1 used the following parameters: TR/TE = 2000/40ms; spectral bandwidth = 1340 Hz; 1024 complex points; VOI = 1.2 x 2.0 x 1.2 cm; 150 water-suppressed repetitions; 2 water reference repetitions. Study 2 used the following parameters: TR/TE = 2000/30ms; spectral width = 1340 Hz; 1024 complex points; VOI = 9 x 27 x 9 cm; 168 water-suppressed repetitions; 2 water reference repetitions. Water suppression was performed using the Variable Power and Optimized Relaxations delays (VAPOR) method. We used TE=33ms on one participant in Study 2 to mitigate increased SAR effects when prompted by the scanner system. The automatic initial shim plus manual higher-order shimming was applied as needed to reduce B0 inhomogeneity. A shim achieving a water peak full-width-half-maximum (FWHM) of 20 Hz or less as estimated by the scanner console was considered acceptable. Data were collected from the left hippocampus if an acceptable shim in the right hippocampus could not be achieved. This was done for 8 young adults and 2 older adults from Study 1 and for 2 older adults from Study 2. Following shimming, the PRESS sequence was collected three times consecutively from the same VOI for both studies. Four older adult subjects were excluded for morphologic abnormalities and two subjects were excluded for poor VOI alignment to the hippocampus.

**Figure 1.**
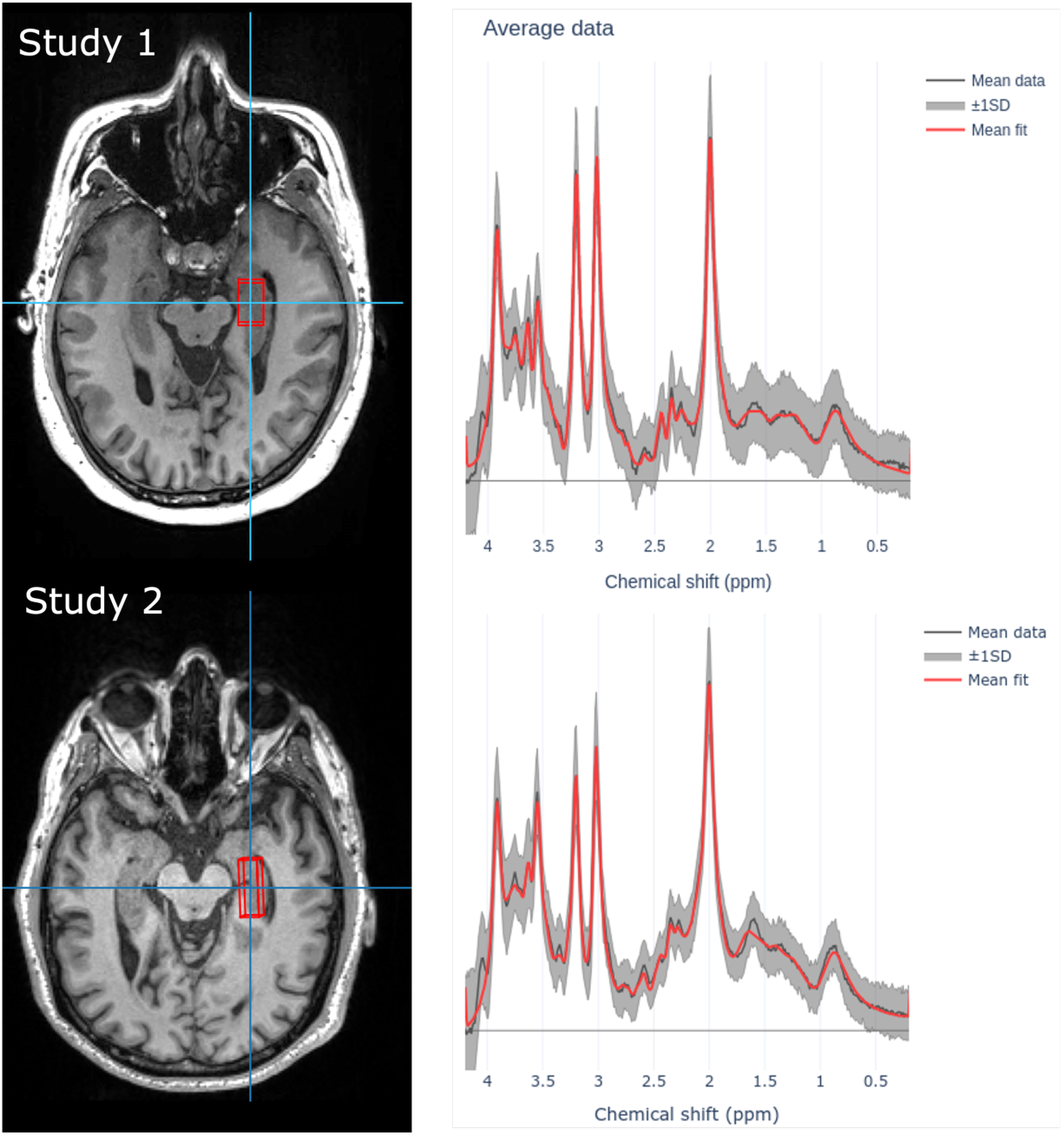
Representative 1H-MRS voxel placement over the right hippocampus for young and older adults in Study 1 and all older adults in Study 2. Mean data (+/− 1SD) and mean fit are shown for each study.

### Data preprocessing

The resulting DICOM-structured data file containing the channel-weighted, preprocessed spectrum was exported and was additionally preprocessed using the open-source Python-based toolbox FSL-MRS (version 2.0.2), part of the FMRIB Software Library (FSL). Expert consensus-recommended practices were implemented^23^ such as eddy current correction, Hankel Singular Value Decomposition (HSVD) water removal, frequency correction, and phase correction^27^. Lastly, FSL-MRS performs fitting of the SVS data using a Linear Combination model where the basis spectra are fitted to the complex-valued spectrum in the frequency domain. Here, we used metabolite basis spectra appropriate for each study’s TE that were simulated externally in TARQUIN^28^. In FSL-MRS, the basis spectra are shifted and broadened with parameters fitted to the data and grouped into 6 metabolite groups. A complex polynomial baseline is also concurrently fitted (order=2). Model fitting was performed using the truncated Newton algorithm as implemented in Scipy. Metabolites were adjusted for tissue volume effects within FSL-MRS which uses FSL’s FAST for tissue segmentation^29^.

Metabolite estimates were considered of initial passing quality with SNR of ≥ 4 and Cramér-Rao lower bound ratios to metabolite concentration (CRLB%) of ≤ 50%. Our key metabolites of interest included MI, tCr, and total NAA (tNAA), which combines NAA and the glutamatergic signaling modulator N-acetyl aspartyl glutamate (NAAG) due to significant overlap of NAA and NAAG at 3T and lower^2^. We initially considered total choline (tCho), as well as glutamate and glutamine (Glx) due to their respective associations with neuroinflammation and excitotoxicity^5^, but these were ultimately removed from further analysis due to estimates below the minimum SNR criterion for nearly half of all subjects. Tukey’s method for extreme outlier detection was used to exclude estimates greater than three times the inter-quartile range for each metabolite.

### Statistical Analysis

Statistical analyses were performed using the statsmodels module (0.14.1) in Python (3.11.8). Robust regressions were used for predicting concentrations for each metabolite using the MacKinnon and White’s heteroscedasticity robust covariance estimator (‘HC2’). Regressions in Study 1 included age group (young vs. old), sex, the FWHM of the spectral linewidth, and signal-to-noise ratio (SNR) as predictors. For Study 2 which was exclusively all older adults, we used the same regressors as Study 1 but replaced the categorical “age group” predictor with actual age in years. Lastly, for the CSF subjects in Study 2, we performed separate robust regressions for each CSF biomarker for predicting concentrations for each metabolite using the same covariates as above, along with using sex- and age-adjusted estimates of ptau181, t-Tau, or Aβ42.

## Results

### Healthy Young versus Older Adult hippocampal metabolite differences

First, Study 1 evaluated metabolite concentrations for MI, tCr, and tNAA in healthy young adults (n=22) ranging from 18 to 32 years old versus older adults (n=18) ranging from 61 to 79 years old to assess typical age-related alterations to hippocampal glial reactivity, bioenergetics, and neural integrity. Robust regressions showed that older adults selectively had greater concentrations of hippocampal MI compared to young adults (p = 0.043, 95% CI [0.069, 4.559]). We also observed milder age-related increases to tCr (p = 0.054, 95% CI [-0.015, 1.925]). No reliable differences were found for tNAA (p = 0.147, 95% CI −0.267, 1.783]) (Figure 2).

**Figure 2.**
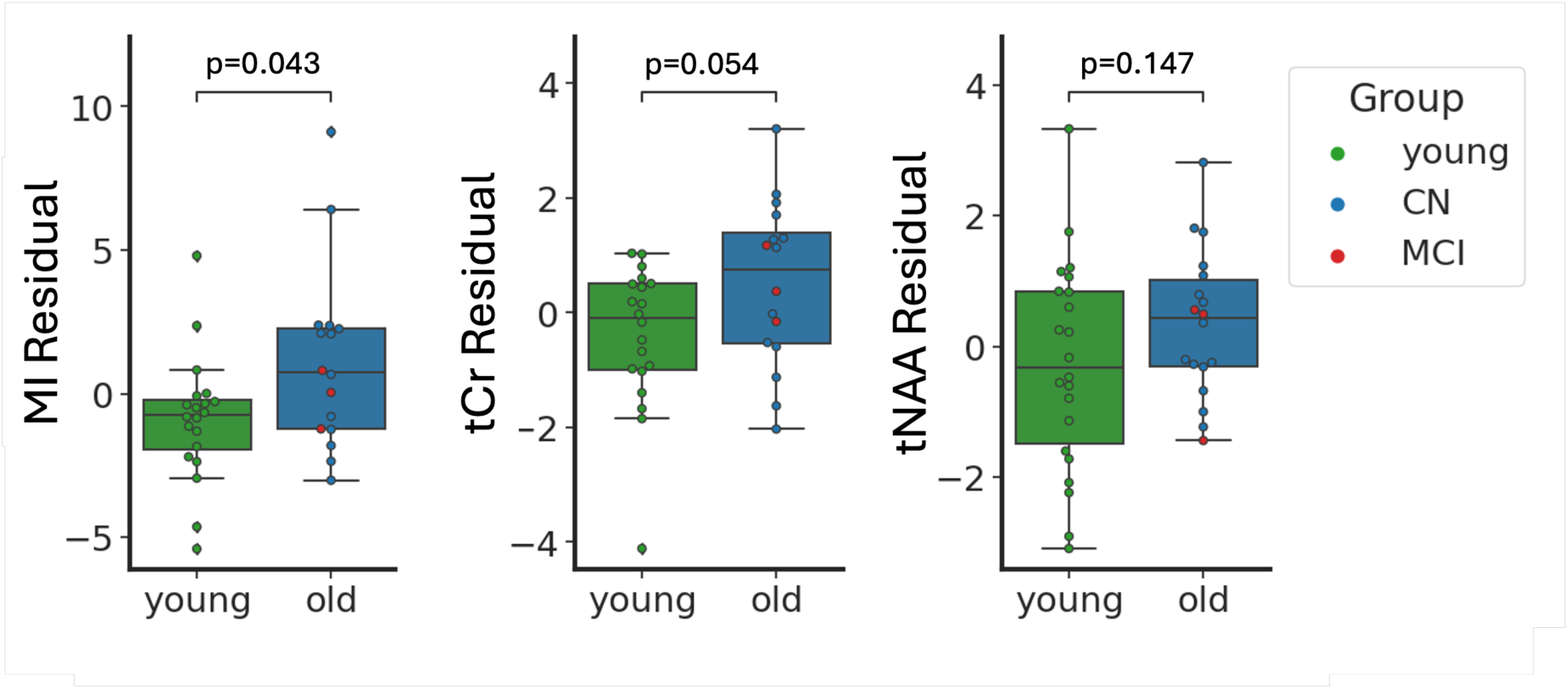
Boxplots of the young versus older adult residuals from robust regression analyses in predicting each metabolite concentration with Sex, FWHM, and SNR as regressed covariates. Subjects with MCI within the older adult group are indicated in red. Older adults showed greater concentrations of MI in the hippocampus compared to young adults (p=0.043). Differences for tCr were trending (p=0.054), and less reliable differences were found for tNAA (p=0.147). CN=cognitively normal; MCI=mild cognitive impairment.

### Total creatine increases as a function of age within older adults

In Study 2, we first examined the effect of age on metabolite concentrations in the hippocampus in a group of cognitively normal older adults (n=29) and an additional six with QCI, MCI, or AD dementia ranging in age from 60 to 85 years. Adjusting for sex, education, FWHM, and SNR, we found that only hippocampal tCr increased with age (p= 0.04, 95% CI [0.004, 0.184]), whereas MI (p= 0.912, 95% CI [-0.142, 0.159]) and tNAA (p= 0.153, 95% CI [-0.024, 0.156]) did not (Figure 3). Notably, only tNAA showed any sex-related differences, where males had higher levels of tNAA compared to females (p=0.008, CI [0.536, 3.482]).

**Figure 3.**
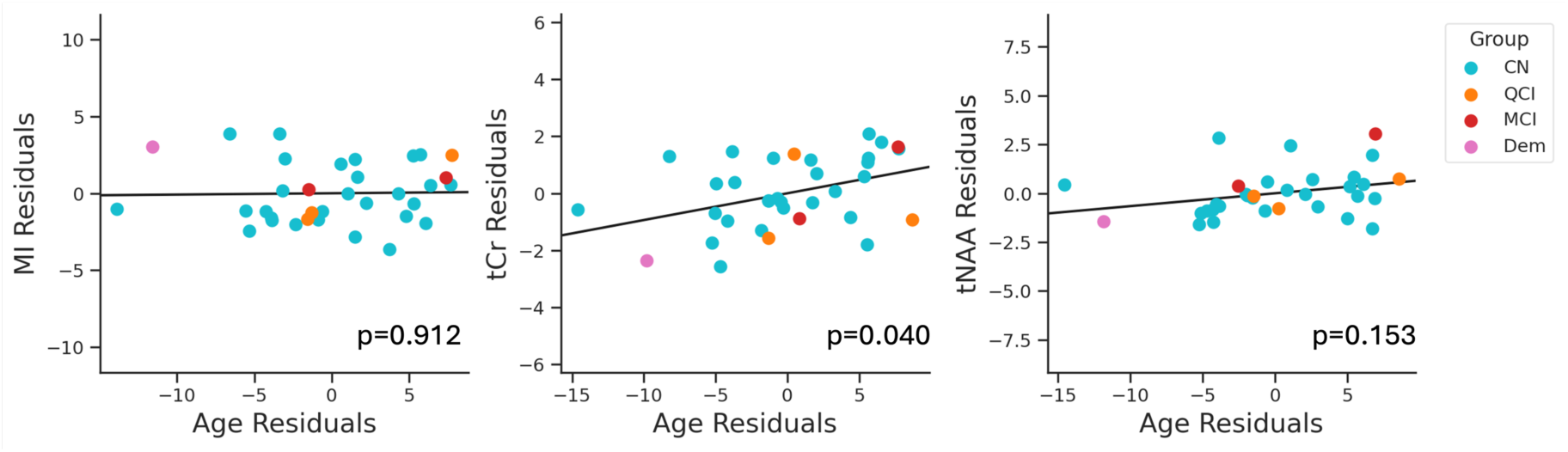
Partial plots of age and metabolites from robust regression analyses predicting each metabolite concentration with Age adjusted for sex, FWHM, and SNR. Older adults showed age-related increases to tCr in the hippocampus. No reliable aging effects were found for MI or tNAA. CN=cognitively normal (blue); QCI=questionable cognitive impairment (orange); MCI=mild cognitive impairment (red); Dem=dementia (pink).

### Metabolites and CSF biomarkers of AD

Finally, we examined relationships between metabolite concentrations in the hippocampus and their relationships to CSF measures of Aβ42, pTau181, and t-Tau in 21 participants from Study 2. Robust regressions showed that adjusted hippocampal MI levels were strongly associated with increasing ptau181 (p<0.001, 95% CI [0.12, 0.331]) and t-Tau (p<0.001, 95% CI [0.017, 0.035]). Importantly, these relationships remained reliable when examining only cognitively normal older adults (ptau181: p=0.0003, 95% CI [0.078, 0.381]; t-Tau: p<0.001, CI [0.013, 0.038]). Furthermore, tNAA was also positively associated with pTau181 (p=0.031, 95% CI [0.013, 0.27]) and t-Tau (p=0.002, 95% CI [0.006, 0.03]). These relationships remained reliable within cognitively normal older adults (ptau181: p=0.016, 95% CI [0.029, 0.282]; t-Tau: p=0.001, 95% CI [0.008, 0.030]). Conversely, no metabolite-CSF relationships were found for tCr or Aβ42 (Figure 4).

**Figure 4.**
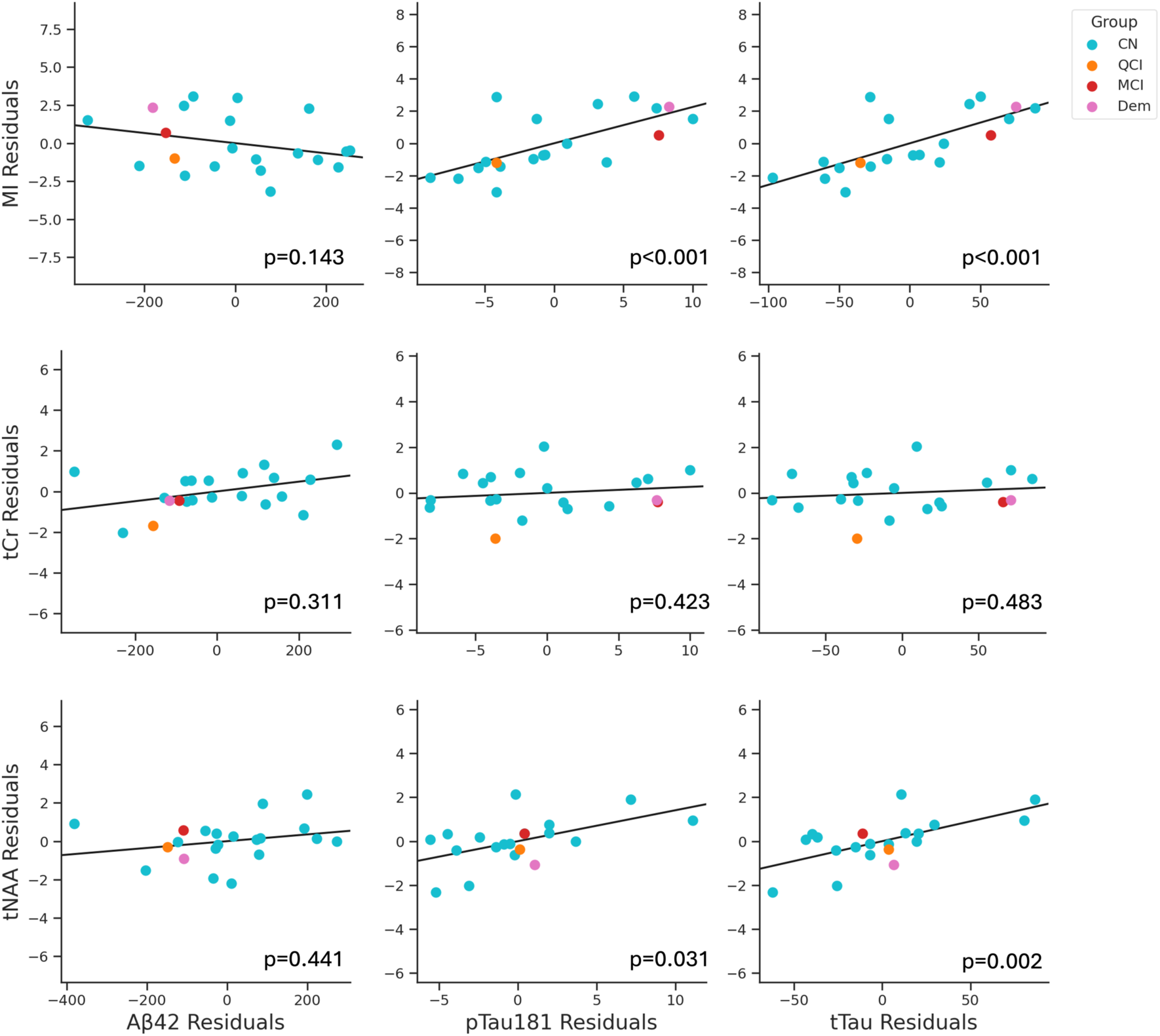
Partial plots and coefficient p-values from robust regression analyses for Aβ42, ptau181, and t-Tau in predicting metabolite concentrations, with age, sex, education, FWHM, and SNR as regressed covariates. MI and tNAA, but not tCr, were linked to increases in CSF pTau181 and t-Tau, but not Aβ42. CN=cognitively normal; QCI=questionable cognitive impairment; MCI=mild cognitive impairment; Dem=AD dementia.

## Discussion

In the present study, we investigated 1H-MRS metabolites in the hippocampus and hypothesized that neurochemical properties associated with neuroinflammation, energy metabolism, and neuronal integrity are associated with increasing age and with CSF AD biomarkers. Our results in cognitively intact young versus older adults identified age-related increases to hippocampal MI and tCr. Additionally, within a separate group of older adults early along the AD continuum, we observed linear age-related increases only to tCr, but not in MI or tNAA. Instead, MI and tNAA were associated with increased CSF ptau181 and t-tau, while CSF Aβ42 was not related to changes to any hippocampal neurometabolites. Overall, our findings indicate that aging and AD exert distinct effects on the neurometabolite profile of the hippocampus, possibly reflecting tau pathology and not Aβ, identifying AD-related changes in the hippocampus before cognitive decline.

Our findings of age-related increases in MI in the hippocampus in older adults is consistent with previous work^3,30^. However, since we did not observe a linear trend between MI and age within older adults and instead showed strong positive associations to CSF ptau181 and t-Tau, this suggests that MI in the older adult hippocampus is more strongly linked to AD pathology rather than increasing age alone. However, few human studies have examined the biochemical properties of the hippocampus in AD, and have primarily focused on AD patients with cognitive impairment and advanced AD pathology with their primary findings linking AD biomarkers to reduced NAA^32,33^. To our knowledge, our findings are the first to begin drawing links between hippocampal neuroinflammation and AD pathology, showing that MI selectively increases with CSF tau pathology even among mostly cognitively normal adults. This suggests that elevated hippocampal neuroinflammation emerges early in the progression of AD prior to overt cognitive impairment and significant neurodegeneration. Mechanistic studies in animal models have artificially increased hippocampal neuroinflammation and found that this to be associated with cognitive deficits ^34,35^. Further, previous studies have shown that increased neuroinflammation is associated with cognitive decline in AD^36,37^ and that heightened glial activation may take place prior to biomarker abnormalities to amyloid and tau^38^. Therefore, understanding the mechanisms that drive hippocampal neuroinflammation, a region with high susceptibility to age- and AD-related dysfunction that is associated with the earliest cognitive symptoms of AD, is essential for developing early interventions and treatments for AD.

Interestingly, we did not observe any relationships with Aβ42 for MI, tCr, or tNAA. This comes in contrast to prior studies that found associations between neurometabolites, Aβ and speed of cognitive decline from MRS collected from early Aβ neocortical sites (i.e., PCC)^12,39,40^. Differences in spatiotemporal spread between amyloid and tau pathologies could potentially explain these differences in outcomes. The pattern of Aβ typically involves early neocortical deposition before spreading to allocortex and subcortical regions^41^. In contrast, tau aggregation early along the AD continuum take place in the hippocampus and entorhinal cortex (Braak stage II), sometimes developing prior to amyloid positivity^42^. Thus, the hippocampal metabolite alterations findings we observed may best reflect early pathological changes related to hyperphosphorylated tau and neurodegeneration, and less with amyloid.

Of note, however, individuals with elevated CSF Aβ42 without tau deposition are at a lower risk for future cognitive decline compared to those with both elevated Aβ42 and tau deposition^43^, demonstrating the synergy of Aβ and tau in giving rise to cognitive impairment. Our current findings begin to offer a mechanistic link for this relationship between tau deposition and cognitive decline, which suggest that hippocampal neuroinflammation may be a key player in this process. However, the present study did not examine relationships to memory performance to test this hypothesis, namely due to the cohort being largely cognitively normal. Future research is needed to determine any direct causal relationships between tau deposition, hippocampal neuroinflammation, and cognitive decline.

Our study using consensus-recommended methods for absolute metabolite quantification did not show any relationships with age and tNAA. Our findings are in line with recent work that used similar methods for examining metabolic changes across the lifespan within the centrum semiovale and posterior cingulate cortex that also found no reliable aging effects on tNAA^44^. Furthermore, the same study found that the FWHM linewidth and SNR of NAA can vary as a function of age and potentially bias overall quantitation and included these quality metrics as covariates in their linear regression analyses. Since the hippocampus can be highly vulnerable to b0 inhomogeneity and reduced SNR, we included age, FWHM, and SNR as covariates as an attempt to account for some of these potential confounding effects. Our findings adjusting for these variables showed that tNAA in the hippocampus is not linked to age but is instead strongly associated with increasing CSF ptau181 and t-Tau.

Although numerous prior studies show age-related decreases to NAA/Cr in typical aging and in AD, and have thus been interpreted as a proxy of neuronal loss and atrophy^3,6^, the presence of alternative mechanisms that alter NAA in the hippocampus remains unclear. A large body of evidence in animal models and human studies have shown that the hippocampus is particularly vulnerable to dysfunctional hyperexcitability in aging and prodromal AD^46–48^, and is a phenotype that may even predict tau accumulation^49^. As NAA is thought to play a role in enhancing mitochondrial energy production from glutamate^45^, we speculate that the novel relationships we observed between increased hippocampal tNAA and CSF ptau181 and with t-Tau may be in part related to increased hippocampal hyperactivity. However, future work examining the basic functional roles of NAA in the aging hippocampus is necessary to better understand the links observed with overall tau pathology.

Lastly, levels of hippocampal tCr, a marker of energy metabolism, were associated with increasing age but not to any CSF AD biomarkers. This demonstrates that not all metabolites within the hippocampus will exert detectable AD-related alterations to the same degree, and that others (i.e., MI and NAA) may be more sensitive to AD pathology and/or pathophysiology.

Furthermore, the positive associations we observed between tCr with age importantly suggest that future studies should exercise caution in interpreting metabolite ratios to Cr especially in context of aging or instead consider performing absolute quantitation of metabolite concentrations.

### Limitations

There was a considerable delay between CSF sample collection and ^1^H-MRS scanning in several participants (average 3 years), with some reaching up to a 7-year delay. If AD pathology has significantly progressed over time, this delay may complicate the interpretations of the relationships between CSF biomarkers and ^1^H-MRS measures. Future work in participants with shorter intervals between CSF and ^1^H-MRS data collection will be useful in further uncovering the relationships between AD pathology and hippocampal biochemistry.

We were unable to evaluate other metabolites of interest such as tCho and Glx due to numerous cases of low SNR and poor fit quality. We speculate that the less than ideal quality of these estimates were due to the known limitations of PRESS within the hippocampus. While other work has shown that results from semiadiabatic LASER sequence (sLASER) and PRESS can be comparable in detecting aging effects in cortical VOIs^50^, the vendor-provided PRESS protocol can be susceptible to magnetic field inhomogeneity in deeper VOIs like the hippocampus (cite). The sLASER sequence has been shown to be robust against these limitations^23,51^. Furthermore, a metabolite-nulled macromolecular spectrum was not collected from each cohort which could aid in reducing model residuals^52^. Future work implementing sLASER and data-driven macromolecule basis sets for hippocampal ^1^H-MRS studies may be better suited for continued assessments of *in vivo* neurometabolic changes in the hippocampus.

A significant strength of ^1^H-MRS is that it can uniquely and non-invasively allow for examinations of specific biochemical changes in the living brain. Yet, it is important to acknowledge that the metabolites measured by this technique are still indirect indicators of the purported functions mentioned (i.e., neuroinflammation, bioenergetics, and neuronal integrity), and since the metabolites measured may participate in more than one physiological process. For example, MI levels have been linked to glial activation and neuroinflammation, but other mechanisms of MI exist, such as in osmolyte regulation and calcium signaling^53^. Whether such mechanisms can also influence overall MI levels independent of neuroinflammation remains to be investigated.

### Conclusions

Our study highlights age-related changes in hippocampal ^1^H-MRS metabolites and their associations with CSF AD biomarkers. We observed increases in MI and tCr with age, while elevated CSF ptau181 and t-tau were associated with higher concentrations of MI and tNAA. These findings suggest that tau pathology, but not Aβ, plays a significant role in hippocampal neuroinflammation and to alterations in neuronal properties even in cognitively normal older adults. Future work examining longitudinal cognitive changes along with ^1^H-MRS can help identify hippocampal metabolites that are predictive of greater risk for future cognitive impairment in AD.

## Acknowledgments

Data collection for Study 1 was funded by NIH R21 AG054092. Funding for the ADRC is supported by NIH P30 AG066519, and support for the research for Study 1 and Study 2 was supported by R01 AG076942. JAN is supported by NIH/NIA T32AG073088, and CV is supported by NIA T35 AG076424. We thank Shirley and Romina for their support in MRI participant recruitment of ADRC subjects, Luke Ehlert for participant recruitment and data collection, Liz Mayer and Hassan Naeem for data collection, and Anna Virovka for data collection and data quality control.

## Notes

### Competing Interest Statement

The authors have declared no competing interest.

